# Functional heterogeneity within the developing zebrafish epicardium

**DOI:** 10.1101/460394

**Authors:** Michael Weinberger, Filipa C. Simões, Roger Patient, Tatjana Sauka-Spengler, Paul R. Riley

## Abstract

The epicardium is essential during cardiac development, homeostasis and repair and yet fundamental insights into its underlying cell biology, notably epicardium formation, lineage heterogeneity and functional cross-talk with other cell types in the heart, are currently lacking. In this study, we investigated epicardial heterogeneity and the functional diversity of discrete epicardial subpopulations in the developing zebrafish heart. Single-cell RNA-sequencing uncovered three epicardial subpopulations with specific genetic programmes and distinctive spatial distribution within the developing heart. Perturbation of unique gene signatures uncovered distinct functions associated with each subpopulation and established novel epicardial roles in cell adhesion, migration, and chemotaxis as a mechanism for recruitment of leukocytes into the heart. This work elucidates the mutual spatiotemporal relationships between different epicardial subpopulations and assigns unique function to each during cardiac development. Understanding which mechanisms cells employ to establish a functional epicardium and to communicate with other cardiovascular cell types during development will bring us closer to repairing cellular relationships that are disrupted during cardiovascular disease.

## Introduction

During embryonic development, numerous distinct cardiovascular cell types closely interact to build and maintain a fully functional heart. The embryonic epicardium is a mesothelial cell sheet that covers the heart’s outer surface and, together with the myocardium and the endocardium, forms the wall of the heart (reviewed in (Simoes and Riley, 2018)). The epicardium derives from a transient extracardiac structure called the proepicardium (Manner et al., 2001). Once the proepicardium envelops the developing heart tube, a subset of epicardial cells undergoes epithelial-to-mesenchymal transition, giving rise to epicardium-derived cells (EPDC) (von Gise and Pu, 2012). These delaminating cells invade the subepicardial compartment and colonise the underlying myocardium to nurture the further growth of the developing heart muscle and coronary vessels, by acting as an essential source of cardiomyocyte and vascular mitogens (Perez-Pomares and de la Pompa, 2011).

In addition to providing signals, EPDCs can directly give rise to many of the key cell types that form the developing heart. Studies have reported an epicardial contribution to adipose tissue (Chau et al., 2014; Liu et al., 2014; Yamaguchi et al., 2015), vascular smooth muscle cells, necessary for vascular support and proper coronary formation, and cardiac fibroblasts in chick (Dettman et al., 1998; Manner, 1999; Mikawa and Gourdie, 1996; Perez-Pomares et al., 1997), mouse (Acharya et al., 2012; Swonger et al., 2016; Wessels et al., 2012; Wu et al., 2013; Zhou et al., 2010) and zebrafish (Kikuchi et al., 2011). A much less consensual view on EPDC fate exists with respect to their putative differentiation into endothelial cells (Dettman et al., 1998; Mikawa and Fischman, 1992; Perez-Pomares et al., 2002; Perez-Pomares et al., 1998; Zhou et al., 2008) and cardiomyocytes (Cai et al., 2008; del Monte et al., 2011; Guadix et al., 2006; Ruiz-Villalba et al., 2013; Zhou et al., 2008). These findings have relied heavily on tissue transplantation and Cre-based fate mapping analyses, which have limitations and may confound interpretation of results due to issues with activation of the Cre drivers in lineage derivatives and mosaic or ectopic expression of reporters (Davis et al., 2012).

While there has been progress understanding the biology of the epicardium and its derivatives, it is still not clear whether pre-migratory EPDCs are a homozygous source of multipotent progenitors or become specified as epicardial subpopulations already within the epicardium proper. To gain an unbiased insight into epicardial cell heterogeneity and the potential functional output of epicardial subpopulations within the embryonic heart, we characterised the developmental transcriptome of the zebrafish epicardium at a single-cell level. We combined confocal microscopy of newly generated epicardial reporter lines and single cell transcriptomics to show the presence of distinct epicardial sub-types in the developing heart. We identified and functionally characterised three transcriptionally distinct epicardial cell subpopulations, only one of which (Epi1) contained cells co-expressing the *bona fide* epicardial signature genes *tcf21*, *tbx18* and *wt1b* (Kikuchi et al., 2011; Serluca, 2008). Functional perturbation using CRISPR/Cas9-mediated gene editing identified *tgm2b*, a transglutaminase gene highly enriched in Epi1, as a new player necessary for the proper development of the epicardial cell sheet enveloping the myocardium. The second subpopulation (Epi2), characterised by significant enrichment in *tbx18*, as well as smooth muscle markers such as *acta2* and *mylka*, was spatially localised outside the main epicardial cell layer, specifically in the smooth muscle covering the outflow tract/bulbus arteriosus (BA) of the developing heart. Loss of the chemokine *semaphorin 3fb* (*sema3fb*), also highly enriched in Epi2, led to an increase in the number of *tbx18* ^+^ cells in the BA, revealing *sema3fb* as an important gatekeeper controlling the spatiotemporal access of epicardial cells to the outflow tract. The third subpopulation (Epi3), that represented a small portion of the *tcf21*-only expressing epicardial cells, was highly enriched for cell migration and guidance cues such as *cxcl12a*. Loss of function of this chemokine resulted in a decreased number of *ptprc/CD45* ^+^ leukocytes recruited to the surface of the epicardium, unravelling the *tcf21* ^+^; *cxcl12a*^+^ Epi3 subpopulation as important in establishing inter-cellular communications responsible for the recruitment and/or retention of *ptprc/CD45* ^+^ haematopoietic cells within the developing heart.

Building on understanding the genetic control of the developing epicardium, our study defines the molecular signatures and novel functional roles of specific epicardial cell subpopulations, thus providing novel insight into the specific programmes within which the epicardium functions during cardiac development.

## Results

### Expression of tcf21, tbx18 and wt1b is heterogeneous in the developing zebrafish epicardium

Expression of the functionally relevant epicardial genes, *Tcf21*, *Wt1* and *Tbx18*, is restricted to subsets of epicardial cells in the developing mouse and chick heart (Braitsch et al., 2012), indicating molecular heterogeneity. Similar findings were observed in the zebrafish epicardium (Gonzalez-Rosa et al., 2012; Kikuchi et al., 2011), however, analysis of expression of the three epicardial genes in the same cell was not previously reported. To gain quantitative insight into the heterogeneity of *tcf21*, *tbx18* and *wt1b* expression within the developing epicardium, we generated the zebrafish triple-reporter line *TgBAC(tcf21:myr-tdTomato;tbx18:myr-eGFP;wt1b:H2B-Dendra2) ^ox187^* by BAC transgenesis (Figs. 1A and 1B). BACs contain extensive genomic fragments, which often include all the regulatory elements required for correct cell type-specific and temporal expression, more closely recapitulating endogenous gene expression patterns than small plasmid-based transgenic lines (Bussmann and Schulte-Merker, 2011). In this triple-reporter line, membrane-tethered tdTomato and eGFP fluorescence label cells expressing *tcf21* and *tbx18*, respectively. Dendra2, localising to the nucleus, identifies cells that express *wt1b*. The resulting fluorescence enabled us to discriminate between epicardial cells that expressed different combinations of *tcf21*, *tbx18* and *wt1b* (Fig. 1C). Confocal microscopy of *TgBAC(tcf21:myr-tdTomato;tbx18:myr-eGFP;wt1b:H2B-Dendra2) ^ox187^* embryos at 3 days post fertilisation (dpf) (Figs. 1D and 1D’), 5dpf (Figs. 1E and 1E’) and 7dpf (Figs. 1F and 1F’) revealed that a large number of epicardial cells did not express *tcf21*, *tbx18* and *wt1b* simultaneously. Close observation revealed cells within the epicardial cell layer covering the ventricle and the outflow tract/bulbus arteriosus (BA) that only expressed different subsets of the three fluorescent markers (Figs. 1D, 1D“’-”’). Quantifying these subsets, we found that the relative amount of *tcf21*, *tbx18* and *wt1b* triple positive epicardial cells increased over time, but it never comprised more than 55% of the fluorescently-labelled epicardial cells (Fig. 1G). To validate the findings obtained using our triple reporter line, we crossed the pre-existing reporter lines *Tg(tcf21:dsRed2) ^pd37^* (Kikuchi et al., 2011) and *Tg(tbx18:dsRed2) ^pd22^*(Kikuchi et al., 2011) to *Tg(wt1b:eGFP) ^li1^* (Perner et al., 2007) and observed clear heterogeneity in the *tcf21* / *wt1b* and *tbx18*/ *wt1b* double-Resuorescent settings (Fig. S1). We also observed epicardial heterogeneity at the endogenous gene expression level by whole-mount in situ hybridisation chain reaction (HCR) (Choi et al., 2010; Choi et al., 2018) (Fig. 1H). At 5dpf, we detected single cell nuclei adjacent to *myl7* positive myocardium that featured endogenous *tcf21* and *tbx18*, but not *wt1b* transcripts (Fig. 1I, asterisk in 1I’). Similarly, we detected *tcf21* and *wt1b*-expressing cells that lacked *tbx18* expression (Fig. 1J, asterisk in 1J’).

**Figure.1.**
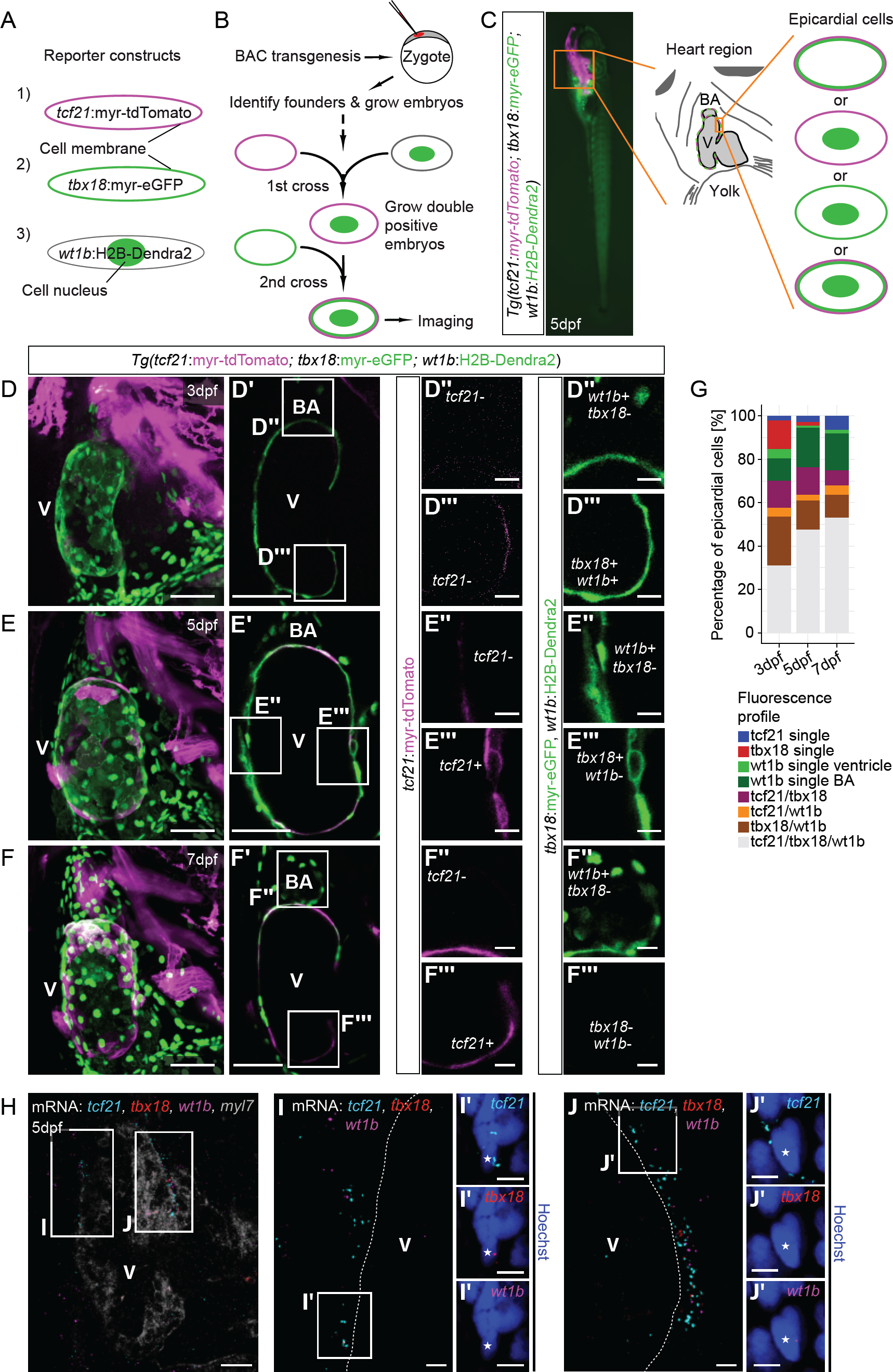
Heterogeneous expression of *tcf21, tbxl8* and *wtlb* in the developing zebrafish epicardium. (A) Subcellular fluorescence of tcf21:myr-tdTomato (magenta membrane), tbx18:myr-eGFP (green membrane) and wt1b:H2B-Dendra2 (green nucleus). **(B)** Workflow to establish the triple reporter line *TgBAC(tcf21:myr-tdTomato; tbx18:myr-eGFP; wt1b:H2B-Dendra2)^ox187^.* **(C)** Schematic of the fluorescence patterns generated by the overlap of tcf21:myr-tdTomato, tbx18:myr-eGFP and wt1b:H2B-Dendra2. **(D-F)** Confocal image projections of the heart region in tcf21:myr-tdTomato, tbx18:myr-eGFP, wt1b:H2B-Dendra2 positive larvae at 3dpf (D), 5dpf (E) and 7dpf (F). **(D‘-F‘)** Single optical sections taken from D-F. **(D“-F“,D“‘-F“‘)** High magnification images of boxed areas in D‘-F‘, depicting representative fluorescence patterns observed in epicardial cells. **(G)** Relative quantification of the different combinations of tcf21:myr-tdTomato, tbx18:myr-eGFP and wt1b:H2B-Dendra2 found in epicardial cells. **(H-J)** Visualisation of *tcf21* (cyan), *tbx18* (red) and *wt1b* (magenta) and *myl7* (grey) mRNA in the larval heart at 5dpf (H). **(I,J)** Magnified images showing the epicardial region (boxed areas in H). **(I‘)** High magnification image of a nucleus (asterisk) in close proximity to *tcf21* and *tbx18,* but not *wt1b* (boxed area in I). **(J‘)** High magnification image of a nucleus (asterisk) in close proximity to *tcf21* and *wt1b,* but not *tbx18* (boxed area in J). Scale bars in D-F,D’-F’: 50 μm, scale bar in H: 20 μm, scale bars in D”-F”,D”’-F”’: 10 pm, scale bars in I,I’,J,J’: 5 μm. V = ventricle, BA = bulbus arteriosus. Number of embryos analysed: 3dpf n=5, 5dpf n=10, 7dpf n=6.

Overall, these results reveal that the expression of *tcf21, tbx18* and *wt1b* in the developing zebrafish epicardium is heterogeneous, both in newly generated BAC transgenic reporter lines and at the level of endogenous gene expression.

### Single cell transcriptomic profiling identifies distinct cell populations within the developing zebrafish epicardium

To further investigate the observed cellular heterogeneity at single-cell resolution, we characterised the epicardial developmental programmes by single-cell RNA sequencing (scRNA-seq) based on the Smart-seq2 technology (Picelli et al., 2013) (see Fig. S2 for quality control data). Since only a small fraction of cells in the heart is epicardial, we FACS-purified cells from hearts extracted from our *TgBAC(tcf21:H2B-Dendra2) ^ox182^*, *TgBAC(tbx18:myr-eGFP) ^ox184^* and *TgBAC(wt1b:H2B-Dendra2) ^ox186^* lines (Fig. 2A). To enhance cell clustering in our data set, we also isolated cells obtained from transgenic hearts where other non-epicardial cardiovascular cells such as cardiomyocytes *(Tg(myl7:eGFP) ^f1^* (Huang et al., 2003), endothelium/endocardium *(Tg(kdrl:GFP)* (Beis et al., 2005) and blood *(Tg(gata1a:dsRed)* (Traver et al., 2003) were labelled, as well as from wild-type hearts. In an attempt to avoid oversampling of *tcf21*-expressing cells, given *tcf21* has previously been described as a pan-epicardial marker (Kikuchi et al., 2011), we also purified non-fluorescent cells from tcf21:dsRed2; myl7:eGFP; kdrl:GFP; gata1a:dsRed quadruple transgenic reporter-positive hearts. Following sequencing, we analysed read count data using the Pagoda pipeline (Fan et al., 2016) to identify cell clusters and sets of divergently-expressed marker genes in an unbiased fashion. To identify distinct cell populations based on shared and unique patterns of gene expression, we applied dimensionality reduction algorithms based on t-SNE, which revealed three distinct epicardial cell populations (Fig. 2B). These populations (termed Epi1, Epi2 and Epi3) were predominantly formed by cells isolated from *TgBAC(tcf21:H2B-Dendra2) ^ox182^*, *TgBAC(tbx18:myr-eGFP) ^ox184^* and *TgBAC(wt1b:H2B-Dendra2) ^ox186^* transgenic hearts, and cells within these subpopulations expressed *tcf21, tbx18* and *wt1b* in a differential manner (Fig. 2C). While *tbx18* was mostly expressed in cells within Epi1 and Epi2, *tcf21* within Epi1 and Epi3, *wt1b* was confined to subsets of cells within Epi1 and, to a lesser extent, Epi3. We further identified 5 distinct non-epicardial cell clusters expressing known markers of other major cell types. These cell clusters comprised cardiomyocytes (Fig. 2C, CM; *vmhc* (Yelon et al., 1999)), erythroid haematopoietic cells (Fig. 2C, eHC; *gata1a* (Lyons et al., 2002)), myeloid/leukocyte haematopoietic cells (Fig. 2C, mHC; *lcp1*(Kell et al., 2018)), neural cells (Fig. 2C, NC; *elavl3* (Park et al., 2000)) and mesenchymal cells (Fig. 2C, MC; *postna*(Snider et al., 2009)). This classification was supported by the enrichment of additional genes that strongly marked each cell population (Fig. S3) and by the over-representation of specific gene ontology (GO) terms within the individual cell populations (Fig. 2D). For example, GO term analysis showed that cells in Epi1, Epi2 and Epi3 expressed many genes associated with epithelial cell fate commitment. Cells in Epi1 were transcriptionally enriched for genes associated with cell adhesion (Fisher’s exact test; p=1.3E-07) and epithelial migration (Fisher’s exact test; p=6.1E-07), fitting the epicardial capacity to migrate as an epithelial cell sheet (Wang et al., 2015). In contrast, cells in Epi2 expressed genes involved in vasoconstriction (Fisher’s exact test; p=0.0227) and cell migration involved in heart development (Fisher’s exact test; p=0.0021), suggesting they might fulfil a function outside the epicardial cell layer. Cells in Epi3 were significantly enriched for genes annotated to white blood cell migration (Fisher’s exact test; p=0.0065) and axon extension (Fisher’s exact test; p=0.0027), suggesting they might be involved in the guidance of non-epicardial cells into the developing heart. To compare the reporter-based results described in Fig. 1 to the endogenous transcriptomic data, we quantified the expression of *tcf21, tbx18* and *wt1b* in Epi1, Epi2 and Epi3 and found that only Epi1 contained cells that co-expressed all three markers (Fig. 2E). Most cells in Epi2 exclusively expressed *tbx18* and many cells in Epi3 exclusively expressed *tcf21.* Collectively our scRNA-seq dataset reveal that the developing zebrafish epicardium at 5dpf consists of three transcriptionally distinct cell populations, with *tcf21, tbx18* and *wt1b* differentially expressed across these populations.

**Figure.2.**
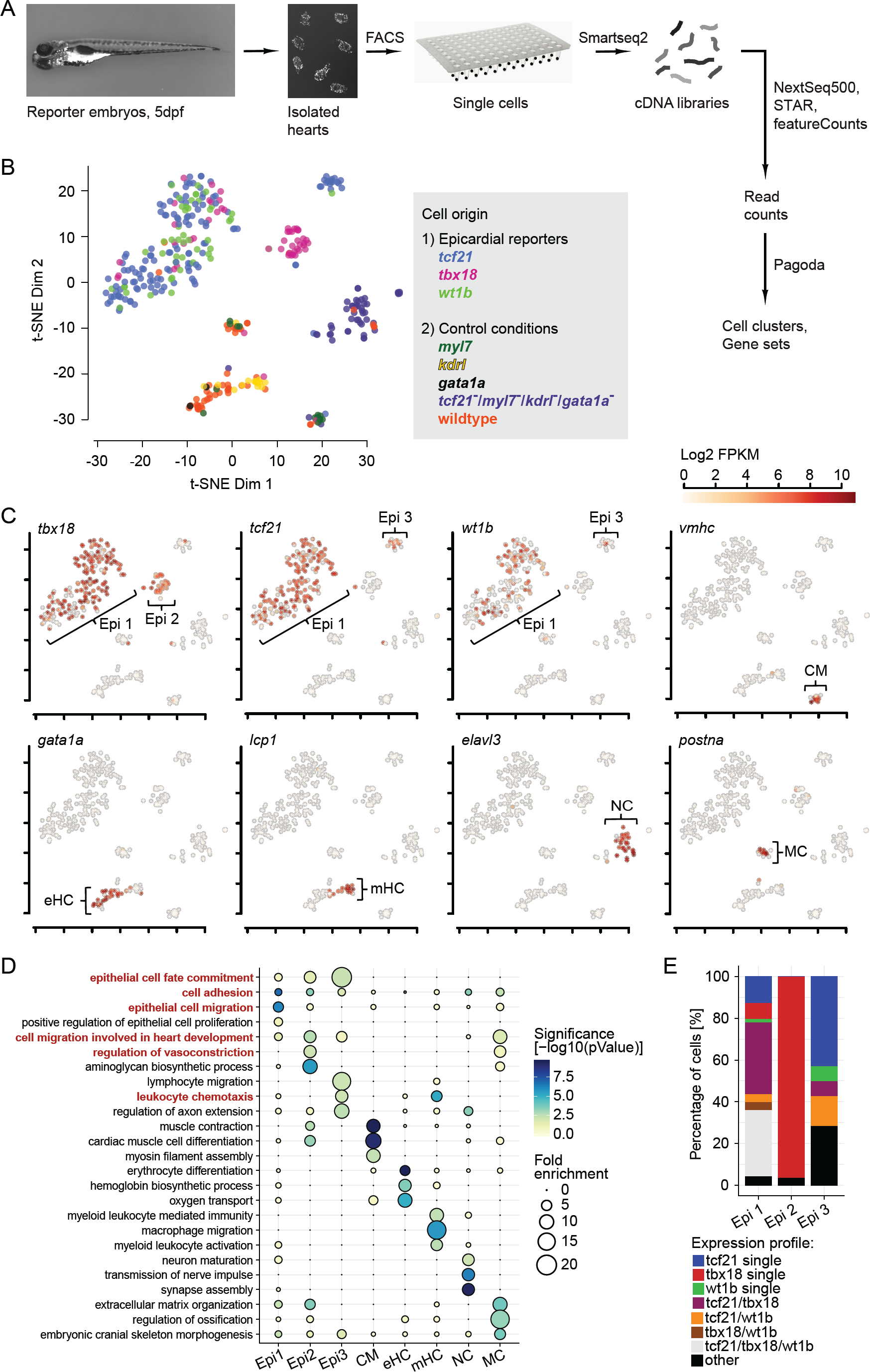
Single cell RNA sequencing reveals distinct epicardial cell clusters in the developing zebrafish heart. (A) Overview of the experimental procedures to prepare, sequence and analyse single cell cDNA libraries from *tcf21, tbx18, wt1b* reporter, control reporter and wildtype hearts. **(B)** t-SNE representation showing clustering in the single cell dataset. Cells are colored according to the reporter/wildtype line they were isolated from. The color key is given in the right-hand legend. **(C)** Identification of single cell clusters based on marker gene expression.The three clusters labelled by *tcf21, tbx18* and *wt1b* are designated Epi1-3. Shown are log2 transformed FPKM values, overlaid onto the t-SNE plot. Expression levels are indicated by the color key. **(D)** GO term over-representation in the different clusters (columns). Bubble size depicts the magnitude of statistical enrichment, the significance is given by the color scale. GO terms important for further analysis are highlighted in red. **(E)** Relative quantification of the different combinations of *tcf21, tbx18* and *wt1b* expression in cells in Epi1-3. CM=cardiomyocytes, eHC=erythroid haematopoietic cells, mHC=myeloid haematopoietic cells, NC=neural cells, MC=mesenchymal cells.

### Epi1, Epi2 and Epi3 are distinct in their transcriptomic profile and spatial distribution within the developing heart

To further investigate the heterogeneity within the epicardial cells in the scRNA-seq dataset, we interrogated the transcriptomic profiles of Epi1, Epi2 and Epi3, which separated into identifiable subpopulations. This analysis uncovered both known and previously uncharacterised genes associated with epicardial development. For example, genes enriched in Epi1 included the retinoic acid synthesising enzyme *retinaldehyde dehydrogenase (aldh1a2),* a known epicardial marker (Kikuchi et al., 2011; Lepilina et al., 2006) (Fig. 3A). However, Epi1 cells also expressed several genes without a previously ascribed epicardial function, such as the signalling peptide *adrenomedullin a* (*adma*), and several adhesion molecules including *junctional adhesion molecule 2b (jam2b)* and *podocalyxin like (podxl),* suggesting that this Epi1 subpopulation might form the main epithelial sheet of cells enveloping the heart. We visualised the expression of *adma* and *jam2b* at 5dpf using multiplexed HCR *in situ* staining and observed that both *adma* and *jam2b* mRNA was present within the epicardial cell layer, co-localised with both *tcf21* and *tbx18* transcripts (Figs. 3B and 3B’) around the same cell nuclei (asterisk in Fig. 3B”). In addition to *tbx18,* Epi2 cells expressed smooth muscle cell markers such as *smooth muscle actin (acta2)* and *myosin light chain kinase (mylka),* as well as extracellular matrix components such as *lysyl oxidase a (loxa)* and *elastin b (elnb)* (Fig. 3A). *elnb* has previously been described as being specifically expressed in the outflow tract/BA of the zebrafish heart (Moriyama et al., 2016). Accordingly, we found the BA to be the only tissue in the developing zebrafish heart at 5dpf that expressed *elnb* mRNA, as well as Mylka protein (Figs. 3C and 3D). Furthermore, *tbx18* mRNA co-localised with *elnb* mRNA in the BA, and multiple Mylka positive cells were labelled by *tbx18* driven myr-Citrine in the same outflow structure. These data suggest that the Epi2 cluster represents an epicardial subpopulation of *tbx18+* cells exclusively located in the smooth muscle layer of the outflow tract/BA. To establish a lineage relationship between Epi2 and the epicardium, beyond the correlation with *tbx18* expression, we performed a tracing experiment using a tcf21-driven Cre *Tg(tcf21:CreER) ^pd42^* (Kikuchi et al., 2011) crossed to the ubiquitous labelling line *Tg(ubi:Switch)* (Mosimann et al., 2011). Similar to previous findings in the adult zebrafish BA (Kikuchi et al., 2011), we observed *tcf21*-derived cells in the smooth muscle layer of the larval BA at 5dpf (Figs. S4A and S4B), supporting an epicardial origin of Epi2 cells rather than an upregulation of *tbx18* mRNA in the outflow tract. However, further lineage tracing with a newly generated *TgBAC(wt1b:Cre-2A-mCherry) ^ox142^* line did not identify any wt1b-derived cells in the BA (Figs. S4C and S4D), indicating that cells in Epi2 never express this transcription factor. RNA velocity analysis of our scRNA-seq dataset, which predictes the future state of individual cells based on distinguishing unspliced from spliced mRNA forms (La Manno et al., 2018), supported the hypothesis that Epi2 cells originate from cells resembling an Epi1 subpopulation (Fig. S5A). Furthermore, pseudo-time trajectories (Trapnell et al., 2014) clearly separated Epi1 from Epi2, but also revealed a subset of cells in Epi1 positioned proximally to Epi2 (Fig. S5B), suggesting transcriptional transition from Epi1 to Epi2 could drive cellular transdifferentiation. This transition state lacks the expression of the smooth muscle marker *mylka,* suggesting migration into the BA precedes the differentiation into a smooth muscle fate (Fig. S5C). Cells in Epi3 strongly expressed *claudin 11a (cldn11a),* the gene coding for an essential tight junction component (Gow et al., 1999) (Fig. 3A). Visualising *cldn11a* mRNA expression, we observed a limited number of positive epicardial nuclei identified by *tcf21* transcripts (Fig. 3E). Most of these nuclei were located in the epicardial region between the BA and the atrioventricular boundary. Therefore, the expression of *cldn11a* was more spatially restricted than the pan-distributed *adma* and *jam2b,* consistent with the small number of cells forming Epi3 in the scRNA-seq dataset (Fig. 2C). Additionally, cells in Epi3 expressed retinoic acid responsive factors such as *nr1h5* and *rxraa,* known to interact in mammalian cells (Otte et al., 2003), as well as the chemokine molecule *cxcl12a* (Fig. 3A). Therefore, this small subpopulation of cells might not only be responsive to retinoic acid but also be involved in the guidance of non-epicardial cells into the developing heart.

**Figure.3.**
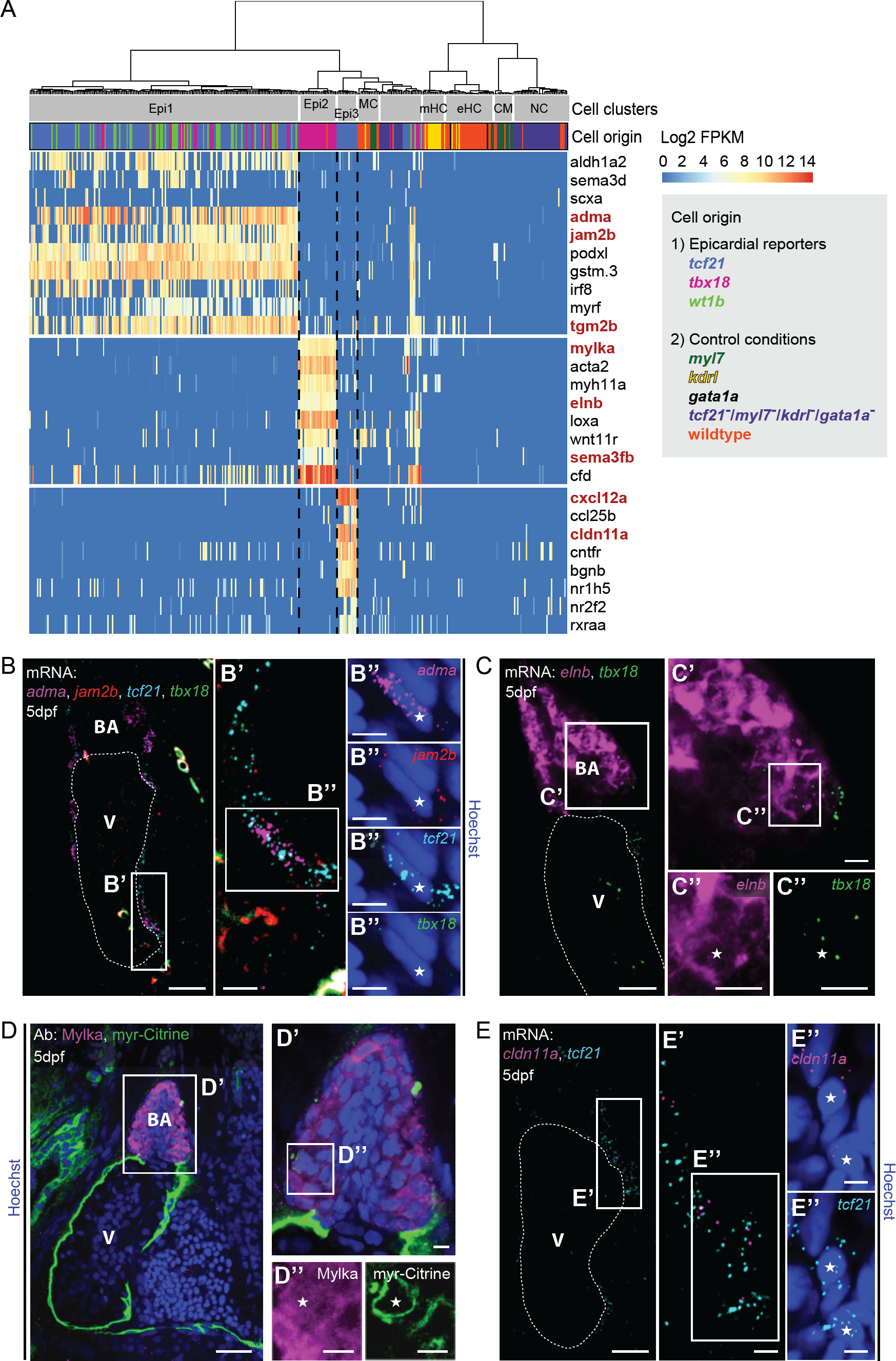
Transcriptionally distinct epicardial subpopulations Epi1-3 localise to different regions of the developing heart. (A)Marker gene expression in Epi1-3. Cells were clustered in an unsupervised manner (columns). Shown are log2 transformed FPKM values. Expression levels are indicated by the color key. Genes analysed further are highlighted in red. Cell cluster identity and color-coded line origin of each cell are indicated at the top of the heatmap. The key is provided in the right-hand box. Black dashed lines indicate the cluster boundaries of Epi1, Epi2 and Epi3. CM=cardiomyocytes, eHC=erythroid haematopoietic cells, mHC=myeloid haematopoietic cells, NC=neural cells, MC=mesenchymal cells. **(B)** mRNA staining of the Epi1 markers *adma* (magenta) and *jam2b* (red), as well as *tcf21* (cyan) and *tbx18* (green) in a 5dpf heart. **(B‘)** Magnified image of the epicardial region (boxed area in B). (B“) High magnification images showing a nucleus (asterisk) surrounded by *adma, jam2b, tcf21*and *tbx18* (boxed area in B‘). **(C)** mRNA staining of the Epi2 marker *elnb* (magenta), as well as *tbx18* (green) in a 5dpf heart. **(C‘)** Magnified image of the BA (boxed area in C). **(C“)** High magnification images showing overlap of *elnb* and *tbx18* (boxed area in C‘). **(D)** Antibody staining of the Epi2 marker Mylka, as well as *tbx18* driven myr-Citrine in a5dpf heart. **(D‘)** Magnified image of the BA (boxed area in D). **(D“)** High magnification images showing overlap of Mylka and myr-Citrine (boxed area in D‘). **(E)** mRNA staining of the Epi3 marker *cldn11a* (magenta), as well as *tcf21*(cyan) in a 5dpf heart. **(E‘)** Magnified image of the epicardial region between BA and atrium (boxed area in E). **(E“)** High magnification images showing two nuclei (asterisks) in close proximity to *cldn11a* and *tcf21* (boxed area in E‘). Scale bars B, C, D, E: 20 μm, scale bars in B’-E’, B”-E”: 5 pm. V=ventricle, BA=bulbus arteriosus.

### The novel Epi1-enriched gene transglutaminase 2b is essential for maintaining the integrity of the epicardial cell layer

To address whether epicardial heterogeneity underlies distinct cell fates and/or function, we used CRISPR/ Cas9 mediated gene knockout of identified Epi1, Epi2 and Epi3 signature genes. In order to achieve efficient and rapid somatic loss of function, it is essential to maintain high expression of guide RNAs (sgRNAs), which are rapidly degraded when not incorporated into Cas9 protein (Burger et al., 2016; Hendel et al., 2015). To achieve this, we made use of the Activator (Ac)/Dissociation (Ds) transposable element system (Emelyanov et al., 2006; Mc, 1950) to stably and efficiently express sgRNAs from a newly generated mini-vector, engineered with a zebrafish U6a promoter (Chong-Morrison et al., 2018). This new approach allows for CRISPR/Cas9-mediated generation of somatic mutagenesis in a highly efficient fashion. The low mosaicism obtained by Ac/Ds-mediated transient transgenesis ensures continued and reproducible expression of sequence-specific sgRNAs with minimal toxicity. To test the efficiency of our transient mutagenesis system, we designed sgRNAs targeting *eGFP* and *citrine,* cloned them into the new Ac/Ds U6a minivector and injected *TgBAC(tbx18:myr-Citrine) ^ox185^* and *TgBAC(tcf21:myr-eGFP) ^ox183^* embryos with the corresponding sgRNA target vector, Ac and Cas9 mRNA. We assessed Cas9-targeting/nuclease efficiency at 5dpf (Figs. S6A and S6B). The analysed embryos showed widespread disruption of fluorescence in the heart region (Figs. S6C, S6D and S6E, S6F), indicating the system was capable of robustly and efficiently perturbing *citrine* and *eGFP* expression using somatic CRISPR/Cas9 genome editing.

*Transglutaminase 2b (tgm2b),* a gene enriched and expressed at high levels within Epi1 cell cluster (Fig. 4A), codes for a protein crosslinking enzyme that has been previously studied during bone formation (Deasey et al., 2012). HCR analysis showed that *tgm2b* transcripts were broadly distributed within the epicardium (Fig. 4B) and localised in close proximity to both *tcf21* and *tbx18* mRNAs surrounding single cell epicardial nuclei (Figs. 4B’ and 4B”), thus suggesting tgm2b might have a specific function within the epicardial cell layer. We designed sgRNAs targeting the region coding for the active site of glutamyltransferase activity, cloned them into the Ac/Ds U6a mini-vector and tested their Cas9-targeting/nuclease efficiency (Figs. S6G and S6H). Somatic knockout of *tgm2b* led to several defects in the epithelial sheet of cells covering the heart at 5dpf (Figs. 4C and 4D). These comprised irregularities in the shape of individual epicardial cells (asterisk in Fig. 4D’) and the formation of aggregates of multiple epicardial cells that had partially detached from the ventricular surface (Fig. 4D”, arrows). In severe cases, large regions of the epicardial cell layer were missing, and epicardial cell numbers at 5dpf were significantly reduced in *tgm2b* knockout embryos (Figs. 4E and 4F). In conclusion, *tgm2b* plays a critical role in maintaining the integrity of the forming epicardium and implicates the Epi1 subpopulation as essential in ensuring the formation of a cohesive epithelial sheet of epicardial cells during heart development.

**Figure.4.**
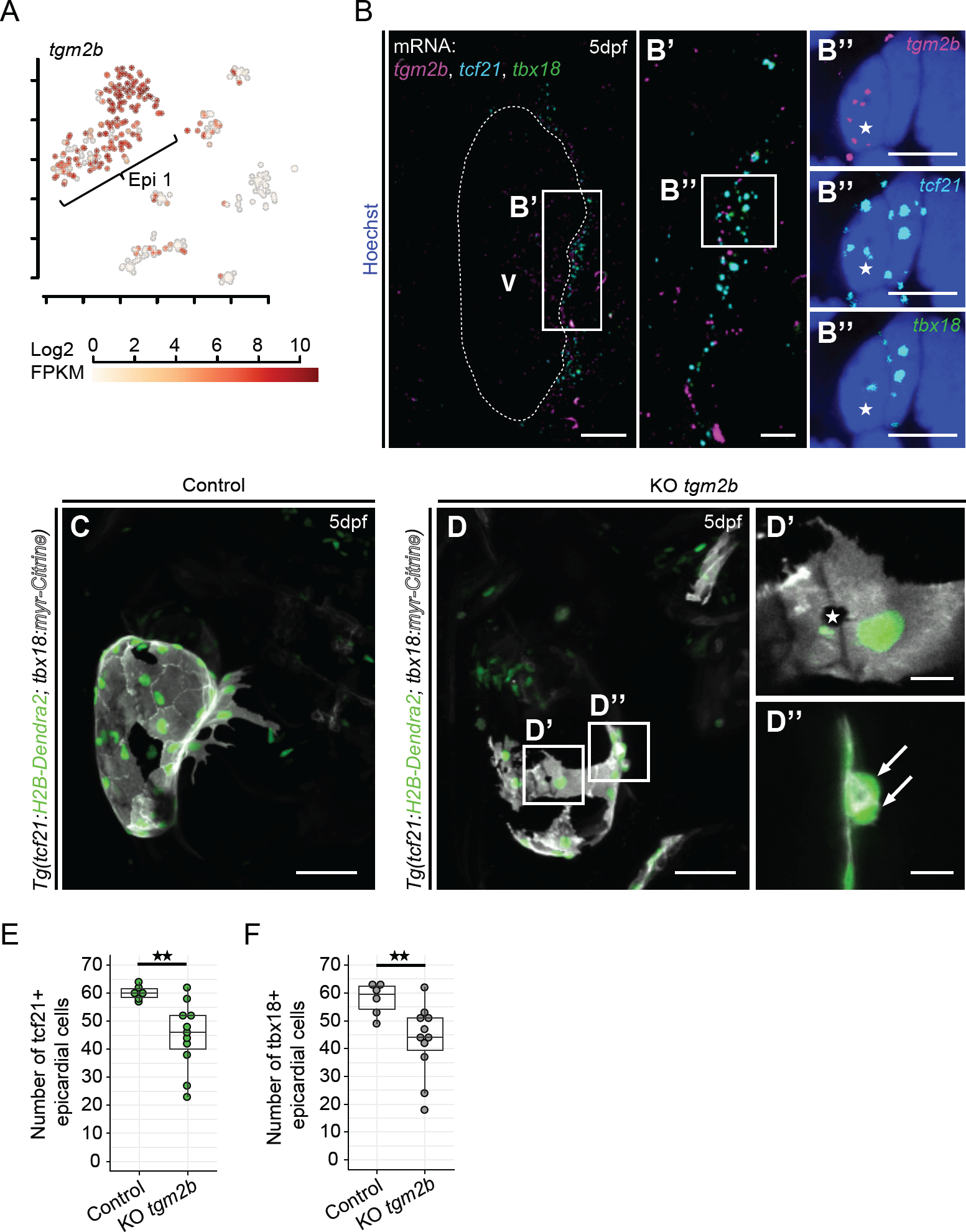
*transglutaminase 2b* is enriched in Epi1 and functions to maintain the integrity of the epicardial cell layer. (A) Expression of *tgm2b*in the scRNA-seq dataset. Shown are log2 transformed FPKM values, overlaid onto the t-SNE plot. Expression levels are indicated by the color key. **(B)** mRNA staining of *tgm2b* (magenta), as well as *tcf21* (cyan) and *tbx18* (green) in a 5dpf heart. **(B‘)** Magnified image of the epicardial region (boxed area in B). **(B“)** High magnification images showing a nucleus (asterisk) surrounded by *tgm2b, tcf21* and *tbx18* (boxed area in B‘). **(C)** The epicardium in a 5dpf control sgRNA injected *tcf21:H2B-Dendra2, tbx18:myr-Citrine* positive larva. **(D)** Disrupted epicardial integrity in a 5dpf *tgm2b* sgRNA injected *tcf21:H2B-Dendra2, tbx18:myr-Citrine* positive larva (KO *tgm2b).* **(D’)** Disruption of epithelial layer integrity (asterisk). **(D”)** Partial detachment of epicardial cells from the ventricular surface (arrows). **(E)** Absolute quantification of *tcf21:H2B-Dendra2*positive epicardial cell numbers in 5dpf control and KO *tgm2b* larvae. **(F)** Absolute quantification of *tbx18:myr-Citrine* positive epicardial cell numbers in 5dpf control and KO *tgm2b* larvae. Scale bars in C,D: 50 μm, scale bar in B: 20 μm, scale bars in D’,D”: 10 pm, scale bars in B’,B”: 5 pm. C and D are confocal image projections. Significance was calculated using Student’s t-test. ** = p<0.01.

### The Epi2-enriched gene sema3fb regulates the number oftbx18+ cells that contribute to the smooth muscle layer of the outflow tract

One of the most enriched factors in the Epi2 cell cluster was *semaphorin 3fb (sema3fb),* a gene coding for a chemo-repellant known to be important for neural patterning (Terriente et al., 2012) (Fig. 5A). Sema3fb belongs to the secreted class 3 semaphorins, which require a receptor complex consisting of a neuropilin (Nrp1a/b or Nrp2a/b) and a plexin for activity (reviewed in (Neufeld et al., 2016)). We surveyed expression of the *sema3* receptors *nrp1a* and *nrp2a* in our scRNA-seq dataset and noticed that *nrp1a* was co-expressed with *sema3fb* in the Epi2 subpopulation, while most of the nrp2a-expressing cells were assigned to the Epi1 cluster, that also featured *nrp1a* expression (Figs. 5B and 5C); suggesting a ligand-receptor intercellular communication within the epicardium. HCR analysis revealed that *sema3fb* was strongly expressed in the BA and only sparsely in the surrounding areas (Figs. 5D and 5D’). In line with our scRNA-seq data, *nrpla* mRNA was present in close proximity to *sema3fb* within the BA, but also in areas adjacent to the outflow tract (Fig. 5D, arrows). Expression of *nrp2a,* on the other hand, was more abundant outside the BA than within (Fig. 5D, arrowheads). However, both *nrpla* and *nrp2a* were expressed at the boundary of the BA, in close proximity to *tbx18* (Fig. 5D’, asterisk) and *tcf21* transcripts (Fig. 5E, asterisks in 5E’, E”), suggesting that both *sema3*cognate receptors are expressed in epicardial cells covering and in close proximity to the outflow tract.

**Figure.5.**
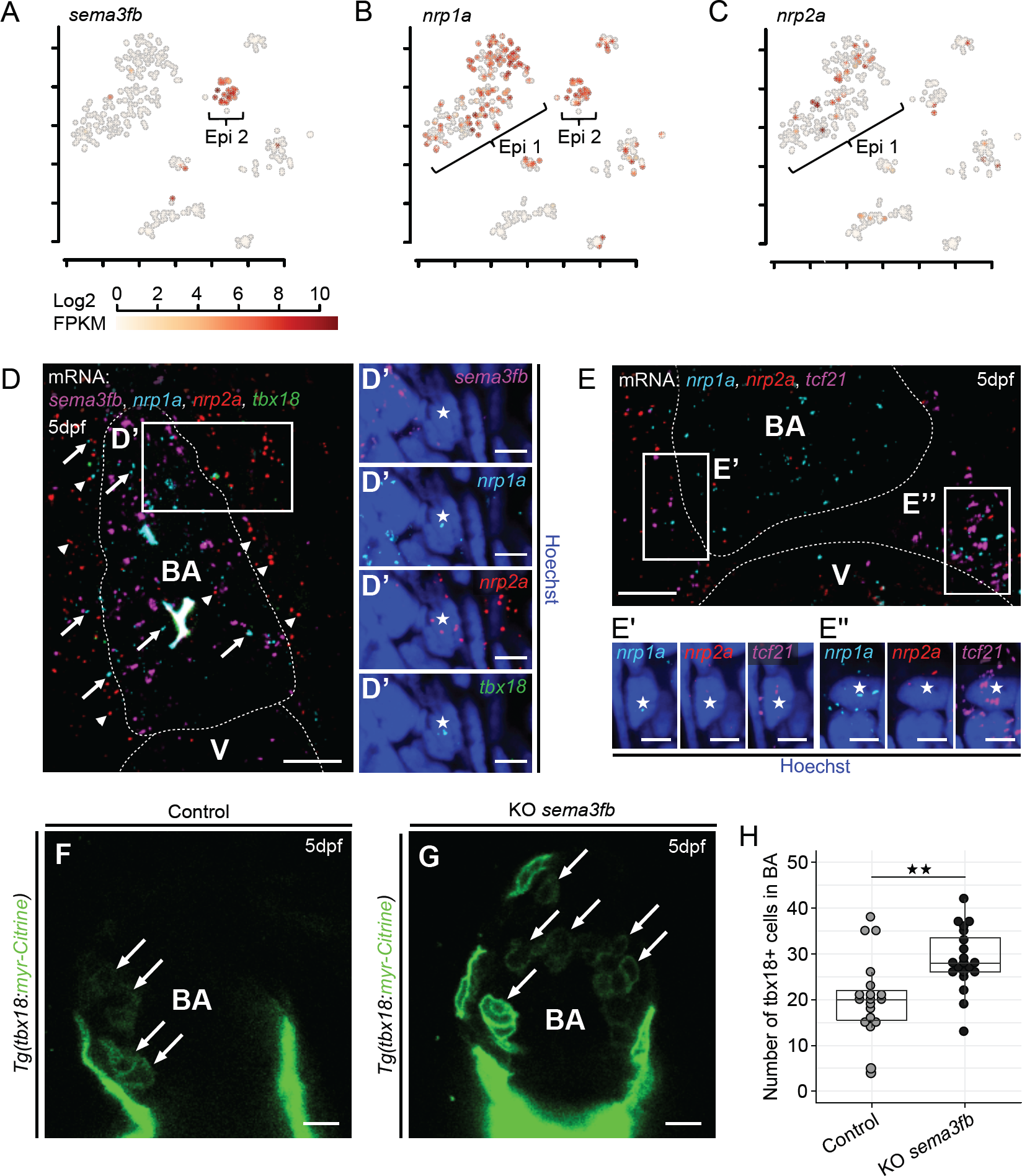
*semaphorin 3fb* is a marker of Epi2 and controls the number of *tbx18+* cells in the bulbus arteriosus. (A) Expression of *sema3fb* in the scRNA-seq dataset. Shown are log2 transformed FPKM values, overlaid onto the t-SNE plot. Expression levels are indicated by the color key. **(B)** Expression of *nrp1a* in the scRNA-seq dataset. **(C)** Expression of *nrp2a* in the scRNA-seq dataset. **(D)** mRNA staining of *sema3fb* (magenta), *nrp1a*(cyan), *nrp2a* (red) and *tbx18* (green) in the BA region of a 5dpf heart. Representative *nrp1a* staining is highlighted by arrows, *nrp2a* is highlighted by arrowheads. **(D‘)** High magnification images showing *sema3fb* in the BA and a nucleus (asterisk) at the BA boundary in close proximity to *nrp1a, nrp2a* and *tbx18* (boxed area in D). **(E)** mRNA staining of *nrp1a* (cyan), *nrp2a* (red) and *tcf21* (magenta) in the region between BA and ventricle at 5dpf. **(E‘,E“)** High magnification images showing nuclei (asterisks) in close proximity to *nrp1a, nrp2a* and *tcf21* (boxed areas in E). **(F)** The BA in a 5dpf control sgRNA injected *tbx18:myr-Citrine* positive larva. **(G)** Increased numbers of *tbx18:myr-Citrine* positive cells in the BA (arrows) in a 5dpf *sema3fb* sgRNA injected *tbx18:myr-Citrine* positive larva (KO *sema3fb).* **(H)** Absolute quantification of *tbx18:myr-Citrine* positive cell numbers within the BA in control and KO *sema3fb* larvae at 5dpf. Scale bars in D,E,F,G: 10μm, scale bars in D’,E’,E”: 5pm. Significance was calculated using Student’s t-test. ** = p<0.01. V=ventricle, BA=bulbus arteriosus.

The expression pattern of *sema3fb* and the *nrp* receptors, together with the significant association of the Epi2 cell cluster to cardiac cell migration (Fisher’s exact test; p=0.00211) (Fig. 2D), suggested that this epicardial subpopulation could be a source of repulsive guidance cues for other epicardial cells migrating into the BA. To test this hypothesis, we designed sgRNAs targeting *sema3fb,* cloned them into the Ac/Ds U6a mini-vector, and tested them for efficient targeting of Cas9-mediated nuclease activity (Figs. S7A and S7B). Somatic mutagenesis of the *sema3fb* locus led to a significant increase in the number of *tbx18+* cells in the BA (Figs. 5F and 5G, quantified in 5H), supporting a role for *sema3fb* in restricting the number of Epi2 cells in the outflow tract. We tested whether this function might be mediated by inhibition of *tbx18+* cell proliferation within the BA (Figs. S7C and S7D, arrows). Control sgRNA and *sema3fb*sgRNA injected embryos were incubated with EdU from 102hpf to 120hpf, after which the number of *EdU;tbx18:myr-Citrine* double-positive cells was quantified. However, we did not detect any significant difference in the number of proliferating *tbx18+* cells in the BA of *sema3fb* knockout hearts as compared to stage-matched controls, despite the fact that many more *tbx18:myr-Citrine* cells were detected within the BA in *sema3fb* mutants (Figs. S7C and S7D, arrowheads and quantification in Fig. S7E). This result argues against a role for *sema3fb* in restricting cell proliferation and suggests that it functions to controls the number of *tbx18+* cells within the BA by restricting their migration from surrounding tissues such as the epicardium.

### Epi3 cxcl12a attracts leukocytes to the developing heart

scRNA-seq defined Epi3 cell cluster was characterized by expression of the chemokine *cxcl12a* (Fig. 6A). HCR revealed that the spatial distribution of *cxcl12a* transcripts was restricted to an area of the epicardial sheet localised between the BA and the atrium (Figs. 6B-6B”). This pattern of expression was very similar to that observed for another Epi3-enriched gene *cldn11a* (Fig. 3E). A subset of the *cxcl12a* mRNA was located in close proximity to *tcf21*transcripts, surrounding single cell epicardial nuclei (asterisk in Fig. 6B”), further validating their co-expression at the single cell level.

**Figure.6.**
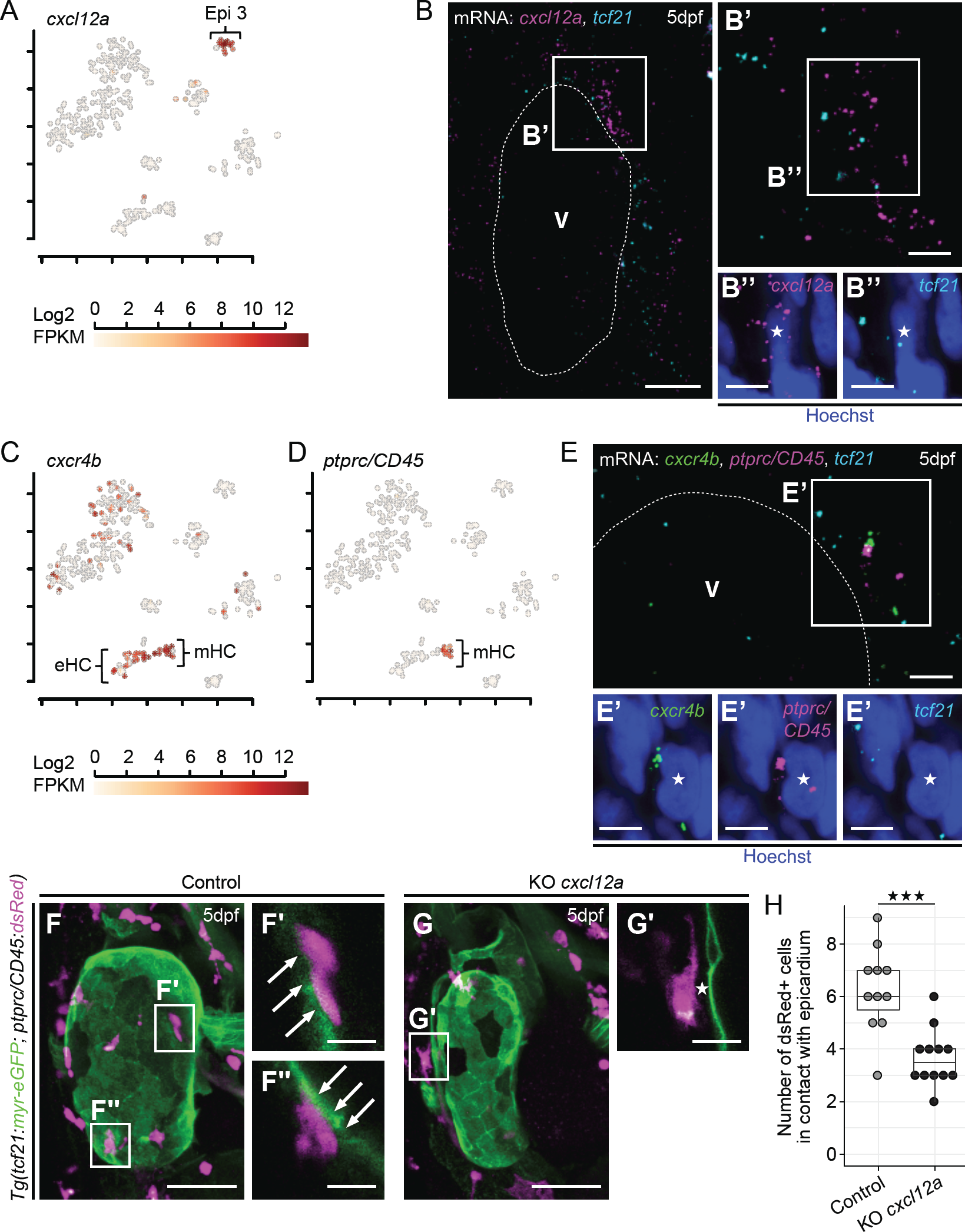
The chemokine *cxcl12a* is expressed in Epi3 and attracts *ptprc+* leukocytes into the epicardium. (A) Expression of *cxcl12a* in the scRNA-seq dataset. Shown are log2 transformed FPKM values, overlaid onto the t-SNE plot. Expression levels are indicated by the color key. (B) mRNA staining of *cxcl12a* (magenta), as well as *tcf21* (cyan) in a 5dpf heart. **(B‘)** Magnified image of the epicardial region between BA and atrium (boxed area in B). **(B“)** High magnification images showing a nucleus (asterisk) surrounded by *cxcl12a* and *tcf21* (boxed area in B‘). (C) Expression of *cxcr4b* in the scRNA-seq dataset. **(D)** Expression of *ptprc* in the scRNA-seq dataset. **(E)** mRNA staining of *cxcr4b* (green), *ptprc* (magenta) and *tcf21* (cyan) in the epicardial region between BA and atrium at 5dpf. **(E‘)** High magnification images showing a nucleus (asterisk) in close proximity to *cxcr4b* and *ptprc* (boxed area in E). **(F)** The epicardium and myeloid blood cells in a 5dpf control sgRNA injected *tcf21:myr-eGFP, ptprc/CD45:dsRed* positive larva. **(F’,F”)** *ptprc/Cd45:dsRed* positive cells are in direct contact with the epicardium (arrows). **(G)** Decreased numbers of *ptprc/CD45:dsRed* positive cells close to the heart in a 5dpf *cxcl12a* sgRNA injected *tcf21:myr-eGFP, ptprc/CD45:dsRed*positive larva (KO *cxcl12a).* **(G’)** A *ptprc/CD45:dsRed* positive cell is located close to the epicardium, but not in direct contact (asterisk). (H) Absolute quantification of *ptprc/CD45:dsRed* positive cells that are in direct contact with the epicardium in 5dpf control and KO *cxcl12a* larvae. eHC=erythroid haematopoietic cells, mHC=myeloid haematopoietic cells. Scale bars in F,G: 50pm, scale bar in B: 20pm, scale bars in F’,F”,G’: 10pm, scale bars in B’,B”,E,E’: 5pm. F,G are confocal image projections. Significance was calculated using Student’s t-test. *** = p<0.001. V=ventricle.

The Epi3 subpopulation was enriched for genes involved in leukocyte chemotaxis by GO term (Fisher’s exact test; p=0.0067), a process that is also significantly associated with the myeloid haematopoietic cell cluster (Fisher’s exact test; p=7.2E-06) (Fig. 2D). Interestingly, analysis of the white blood cell (mHC) transcriptome (Fig. 3A) revealed that the *cxcl12a* cognate receptor *cxcr4b* was mostly present in this population (Fig. 6C, mHC), which is also unique for its expression of *ptprc (CD45)* (Fig. 6D), a pan-leukocyte marker in zebrafish (Bertrand et al., 2008). We validated these findings by HCR and found distinct cell populations in which the transcripts of *cxcr4b* receptor and *ptprc/CD45* gene were co-localised (Fig. 6E). In some cases, this was evident at the ventricular surface (Fig. 6E’, asterisk), with the expression of *tcf21* in the neghbouring cells, but never overlapping with either *cxcr4b* or *ptprc/CD45* transcripts. This suggested that epicardial-derived *cxcl12a* chemokine might be attracting *ptprc/CD45+* leukocytes to the developing heart. To test this hypothesis, we designed sgRNAs targeting the *cxcl12a* gene and cloned them into the Ac/Ds U6a mini-vector. We verified sgRNA Cas9-mediated nuclease activity efficiency using our restriction assay (Figs. S7F and S7G) and targeted the *cxcl12a* locus in embryos obtained from crossing the newly generated *TgBAC(tcf21:myr-eGFP) ^ox183^* to the leukocyte reporter line *Tg(ptprc/CD45:dsRed) ^sd3^* (Bertrand et al., 2008). We found that somatic loss of *cxcl12a* decreased the number of ptprc/CD45:dsRed+ cells that were recruited into and/or retained on the surface of the heart (Figs. 6F and 6G, quantified in 6H). Additionally, while we observed direct contact between *ptprc/CD45:dsRed+* cells and the epicardium in control embryos (arrows in Figs. 6F’ and 6F”) this appeared to be disrupted in the *cxcl12a* knockout background (Fig. 6G’, asterisk). This data identifies a novel role for the Epi3-derived *cxcl12a* in chemoattracting leukocytes to the surface of the developing heart.

## Discussion

Understanding epicardial cell fate and lineage potential to-date has been constrained by the limited set of so-called canonical markers, which are either not entirely specific for epicardial cells or fail to label the entire epicardial cell pool. Here we identify epicardial cell subpopulations within the developing zebrafish heart, characterise their transcriptional profiles at the single-cell level and functionally test their involvement during heart development in zebrafish.

Our study provides unique insights into the heterogeneity and associated functional diversity of the developing epicardium, unravelling novel epicardial functions that go beyond previously characterised mitogenic roles and known contribution of cardiovascular derivatives during development (Perez-Pomares and de la Pompa, 2011). We reveal the distinct genetic programmes, and their associated spatial distributions, harboured within newly-identified Epi1, Epi2 and Epi3 epicardial cell subpopulations/clusters (Fig. 7A). Gene ontology analysis of the most highly enriched genes expressed in each of the three epicardial cell clusters and CRISPR/Cas experimental perturbation of signature genes provided novel insights into the maintenance of epicardial integrity, formation of the outflow tract and haemaopoeitic cell cross talk and recruitment into the developing heart.

### Maintenance of epithelial integrity within the forming epicardium

Epi1-expressed genes were significantly enriched for GO terms related to biological processes of cell adhesion and epithelial migration, consistent with a possible role in the formation of a coherent epicardial cell sheet that migrates to envelop the myocardium. We identified a novel epicardial gene, *tgm2b,* which when functionally perturbed, resulted in disruption of the integrity of the epicardial epithelium and impaired epicardium formation overlying the myocardium (Fig. 7B). The reduced number of epicardial cells that envelop the *tgm2b*-mutated heart might be due to a reduction in the number of epicardial cells that migrate from the PE and attach to the myocardium (Hatcher et al., 2004; Hirose et al., 2006; Ishii et al., 2010). Alternatively, tgm2b could be required to modulate the epicardial-myocardial interactions required during the actual assembly of the epicardium onto the heart (Kwee et al., 1995; Sengbusch et al., 2002; Yang et al., 1995), a process also known to require heartbeat-derived fluid flow (Peralta et al., 2013). The appearance of cell aggregates and significant interruptions to the epithelial layer in the mutants suggests tgmb2 acts to ensure cell-cell contact as an essential process during epicardium formation.

### The regulation of epicardial cell migration into the outflow tract

Epi2-enriched genes were significantly associated with the biological processes of cell migration involved in heart development and vasoconstriction, suggesting that this subpopulation might be linked to the smooth muscle wall of the ventricular outflow tract/bulbus arteriosis. Indeed, many Epi2-specific smooth muscle associated genes were exclusively expressed in the developing smooth muscle cell layer surrounding the BA endothelium. We showed that these *tbx18+* Epi2 cells had a *tcf21* origin but never expressed *wt1b.* Pseudo-time analysis suggested that Epi1 cells might be in a transition state towards Epi2, (Fig. S5), and this was confirmed by our *tcf21* and *wt1b* lineage tracing results. The outflow tract of the adult zebrafish heart has been identified as a positive regulator of epicardial cell migration (Wang et al., 2015). Following cardiac injury, the BA is thought to drive the coordinated movement of the regenerating epicardial cell sheet, in a manner that is dependent upon short-range BA-derived Sonic hedgehog signalling (Wang et al., 2015). One of the most enriched genes in the developing Epi2 subpopulation encoded the secreted diffusible chemorepulsive factor sema3fb, which we revealed functions within the BA to negatively regulate the number of epicardial cells that populate the BA (Fig. 7C), therefore, acting as a gatekeeper controlling the appropriate number of cells in the outflow tract. Both sema3 signalling receptors, *nrp1a* and *nrp2a,* were expressed in the epicardium albeit with distinct expression patterns. *Nrp1a* was co-expressed with *sema3fb* in the Epi2/BA subpopulation, while most of *nrp2a* transcripts were confined to the Epi1/epithelial sheet covering the heart, which also expressed *nrp1a.* NRP1 and NRP2 have been reported to differentially interact with SEMA3 ligands, with SEMA3F specifically binding to NRP2 and SEMA3A partnering with NRP1 (reviewed in (Neufeld et al., 2016)). The spatial distribution of *nrp1a* and *nrp2a* transcripts in the developing heart could account for the different way the migrating epicardial cells respond to the same sema3fb diffusible cue. Cells in Epi1, which express the sema3fb-responsive receptor nrp2a, are retained outside of the BA by the chemorepulsive activity of sema3fb, while epicardial cells that migrate into the BA only express *nrp1a,* and not *nrp2a*, and are, therefore, able to home from the epicardium to the outflow tract. Our work highlights the importance of understanding how cell-cell interactions are established during the development of the heart and, in particular, the mechanisms enabling migrating epicardial cells to respond differently to the same diffusible chemorepulsive cue.

**Figure.7.**
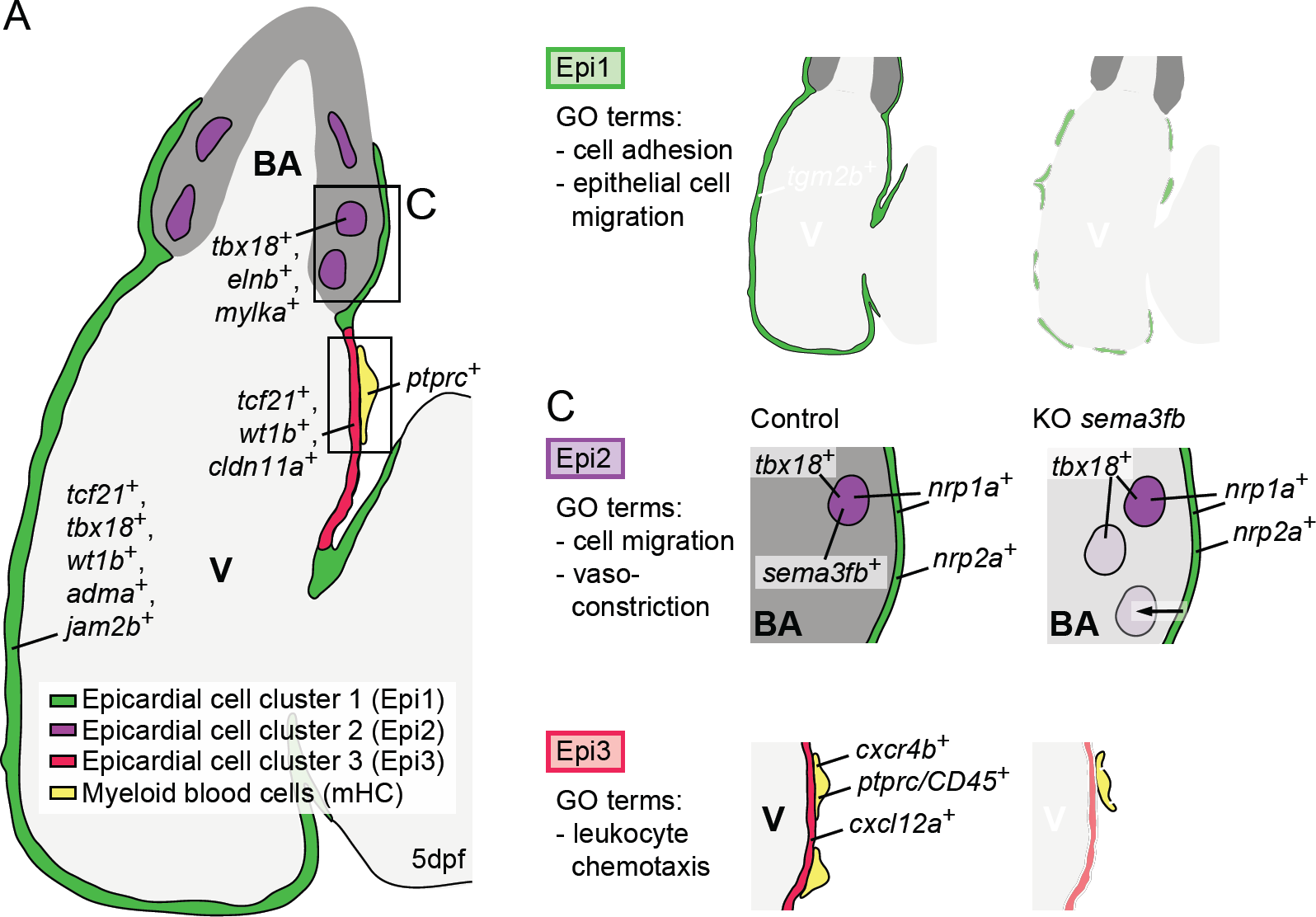
Summary of distinct gene expression, spatial distribution and function of epicardial cell subpopulations in the developing zebrafish heart. (A) Depiction of the zebrafish heart at 5dpf. The locations of Epi1, Epi2 and Epi3 are indicated in color-code. The color key is at the bottom of the schematic. **(B)** Loss of *tgm2b* (KO *tgm2b)* leads to disrupted integrity of the epicardial cell layer. **(C)** Loss of *sema3fb* (KO *sema3fb)* leads to increased numbers of Epi2 cells in the BA, possibly due to increased cell immigration from the epicardium. **(D)** Loss of *cxcl12a* (KO *cxcl12a)* leads to a decreased presence of mHC cells on the heart. V=ventricle, BA=bulbus arteriosus.

### Epicardial cell regulation of leukocyte homing into the heart

Genes in the Epi3 cell cluster were associated with the processes related to leukocyte chemotaxis, indicating that this subpopulation might be involved in the guidance of white blood cells into the developing heart. Links between the epicardium and the immune system have been implicated during the development of the mouse fetal heart, whereby *CD68+* macrophages were shown to require the intact epicardium for their recruitment into the developing heart, in a process that was dependent on the embryonic expression of *Wt1* (Stevens et al., 2016). Our study similarly reveals that the Epi3-enriched chemokine ligand *cxcl12a* is necessary for the recruitment and/or retention of a distinct population of *ptprc/CD45+* haematopoietic cells onto the surface of the heart (Fig. 7D). These leukocyte/myeloid cells express the cognate chemokine receptor *cxcr4b.* Therefore, our work unravels a novel mechanism by which a specific epicardial subpopulation developmentally regulates leukocyte recruitment through the expression of a paracrine signalling molecule. Non-PE derived *CD45+* haematopoietic cells were shown to contribute to the developing epicardium of the mouse heart (Balmer et al., 2014). Therefore, it will be of interest to further explore the possible role of *CD45+;cxcr4b+* cells that migrate onto the developing zebrafish heart. *Cxcl12a* is also expressed in adult epicardial cells upon cardiac injury (Itou et al., 2012) and, in the mouse, epicardial-associated *CD45+* haematopoietic cell clusters respond dynamically to myocardial infarction (Balmer et al., 2014). These cells co-express *CD45+/CXCR4+/CD11b+*in the bone marrow and within the ischemic heart (Ghadge et al., 2017), suggesting that *cxcl12a* may recapitulate its developmental role following adult heart injury.

In conclusion, our scRNA-seq work has opened up new avenues for understanding epicardial cell biology and provides novel insight into formation of a critical lineage which is essential during both heart development and adult heart regeneration. The identification of functional subpopulations of epicardial cells and their roles as sources of chemo-attractant and repellent signals for cellular cross-talk during development may be exploited to facilitate cell-based therapies to regenerate the injured heart.

## Author contributions

Conceptualisation, FCS, TSS and PRR; Methodology, MW and FCS; Investigation, MW and FCS; Writing-Original Draft, MW and FCS; Writing-Review & Editing, TSS and PRR; Bioinformatic analysis and Data Curation, MW; Supervision, FCS, TSS and PRR; Funding Acquisition, RP, TSS and PRR.

## Acknowledgements

We would like to thank Biomedical Services Unit for fish husbandry; the WIMM Flow Cytometry Facility; the WIMM Sequencing Facility and The Wolfson Imaging Centre. We would like to thank Vanessa Chong-Morrison for providing us with the zebrafish U6a promoter Ac/Ds sgRNA expression vector.

*Tg(tcf21:dsRed2) ^pi37^* and *Tg(tbx18:dsRed2) ^pd22^* lines were kindly provided by Ken Poss, *Tg(wt1b:eGFP)^1^’1* by Christoph Englert, *Tg(ptprc/CD45:dsRed) ^sd3^* by David Traver and *Tg(ubi:CSY)* by Marianne Voz.

*Tg(ubi:Switch)* was obtained from the Zebrafish International Resource Center (ZIRC). This work was supported by a British Heart Foundation (BHF) 4 year PhD studentship FS/13/57/30647 to MW, an Oxford BHF Centre of Research Excellence Fellowship RE/13/1/30181 award to FCS; MRC-MHU grant to RP; MRC grant (G0902418), and Oxford BHF CRE award (RE/08/004) to TSS and BHF grants RG/13/9/303269 and CH/11/1/28798 to PRR.

## Experimental Procedures

### Ethics statement

All animal experiments were performed under a Home Office Licence according to the Animals Scientific Procedures Act 1986, UK, and approved by the local ethics committee.

### Zebrafish

Published transgenic reporter lines used in this study were:*Tg(tcf21:isRei2) ^pi37^* (Kikuchi et al., 2011), *Tg(tbx18:dsRed2) ^pd22^* (Kikuchi et al., 2011), *Tg(tcf21:CreER) ^pd42^* (Kikuchi et al., 2011), *Tg(wt1b:eGFP) ^h1^* (Perner et al., 2007), *Tg(myl7:eGFP) ^f1^* (Huang et al., 2003), *Tg(kdrl:GFP) ^s843^* (Beis et al., 2005), *Tg(gata1a: dsRed)*(Traver et al., 2003), *Tg(ptprc/CD45:dsRed) ^sd3^* (Bertrand et al., 2008), *Tg(−3.5ubi:loxP-EGFP-loxP-mCherry) [Tg(ubi:Switch)]* (Mosimann et al., 2011) and *Tg(−3.5ubi:loxPAmCyanSTOPloxPZsYellow) [Tg(ubi:CSY)]* (Zhou et al., 2011). *Tg(tcf21:dsRed2; myl7:eGFP)* and *Tg(kdrl:GFP; gata1a:dsRed)* double transgenic lines were generated by natural mating.

### Generation of transgenic animals

To generate *TgBAC(tcf21:myr-tdTomato)*^ox181^, *TgBAC(tcf21:H2B-Dendra2)^ox182^, TgBAC(tcf21:myr-eGFP)ox183, TgBAC(tbx18:myr-eGFP)*^ox184^, *TgBAC(tbx18:myr-Citrine)*^ox185^, *TgBAC(wt1b:H2B-Dendra2) ^ox186^* and *TgBAC(wt1b:Cre-2A-mCherry) ^ox142^* we used a BAC recombineering approach (Bussmann and Schulte-Merker, 2011). BACs used were DKEYP 79F12 *(tcf21),* DKEYP 117G5 *(tbx18)* and CH73 157N22 *(wt1b).* Dendra2, tdTomato, eGFP and Citrine fluorophore sequences linked to 5’ histone 2b (H2B) or 2x myristoylation (myr) sequences (Dempsey et al., 2012; Zacharias et al., 2002) were inserted into the first coding exon of *tcf21, tbx18*or *wt1b* BACs via bacterial homologous recombination as previously reported (Trinh et al., 2017). Cre-2A-mCherry was inserted similarly into the *wt1b* BAC. The corresponding constructs used for amplification of donor fragments flanked with recombinant arms are available from Addgene. The flanking sequences for homologous recombination were: *tcf21-5’* 5’-CATCTCCTCAAGAAGTCCTTTTCTCCACTCCACCCTTGTCTCCAGCCAAC-3’, *tcf21*-3’ 5’-CCAAACAACATTAGATTAACCGAATCGGAAAACCAAAATGAATTTATGAAAACTCAATATTAATTCTGATTGCAAGAAGTGTCTCAC-3’, *tbx18-5’* 5’-TTCTGGTGAACTTCTCTTTCTCGGCCAATCTGTCTTCTCGGTCGGTAACC-3’, *tbx18-3’* 5’-TACTTACGGGATTCGTCGCTGGTGCAATCTATCTCGCAACTCCTGGTGCT-3’, *wt1b-5’* 5’-GTGTTTTGCAACCCAGAAAATCCGTCTAAATGCTGACAGAGCCGTGCGGCCCG-3’ and wt1b-3’ 5’-GACCACATTGAGAGAGATTTTGAGGCGAGATTGTAAGGACGGGATGGTTTTCTCAC-3’. BACs were injected into one-cell zygotes and integrated into the genome via Tol2-mediated recombination. *TgBAC(tcf21:myr-tdTomato;tbx18:myr-eGFP;wt1b:H2B-Dendra2) ^ox187^* was generated by two consecutive rounds of breeding from *TgBAC(tcf21:myr-tdTomato) ^ox181^*, *TgBAC(wt1b:H2B-Dendra2) ^ox186^* and *TgBAC(tbx18:myr-eGFP)^ox184^.*

### Hybridisation chain reaction

Embryos were fixed in 4% paraformaldehyde (PFA) overnight at 4 °C. Subsequently, embryos were stored at-20 °C in methanol. Hybridisation chain reaction (HCR) v3.0 (Choi et al., 2018) was performed following a protocol by Choi et al. Briefly, larvae were permeabilized using 30 |jg/ml proteinase K for 45 minutes at room temperature, postfixed in 4% PFA and incubated overnight at 37 °C in 30% probe hybridisation buffer containing 2pmol of each probe mixture. Excess probes were washed off with 30% probe wash buffer at 37 °C and 5xSSCT at room temperature and larvae were incubated overnight at room temperature in amplification buffer containing 15pmol of each fluorescently labeled hairpin. Following HCR, larvae were incubated with Hoechst reagent (1:1000, 5xSSCT) for 30 minutes at room temperature. Images were obtained using a LSM780 confocal microscope (ZEISS) and a 40x objective.

### Immunocytochemistry

Larvae were fixed in 4% PFA for 45 minutes at room temperature. After washing, larvae were blocked using 5% goat serum (PBS, 0.5% Triton, 2% DMSO) for 1 hour at room temperature. Primary antibodies used were: chicken anti-GFP (ab13970, Abcam), rabbit anti-dsRed (632496, Clontech) and mouse anti-Mlck (M7905, Sigma-Aldrich) in a 1:500 dilution, added overnight at 4 °C. Secondary antibodies used were: goat anti-chicken/488nm, goat anti-rabbit/555nm, goat anti-mouse/633nm and Hoechst reagent in a 1:1000 dilution, added for 2 hours at room temperature.

### EdU staining

Larvae were incubated in embryo medium containing 1mM EdU and 1% DMSO for 18 hours, from 102hpf to 120hpf. Larvae were stained by immunocytochemistry against GFP, following detection of EdU using the Click-iT Alexa Fluor 647nm kit (ThermoFisher) for 2 hours at room temperature.

### Heart purification and FACS

Embryonic hearts were isolated following a published protocol (Burns and MacRae, 2006), using a 21-gauge needle for disruption. 150-200 hearts were collected from 250-350 larvae. Hearts were dissociated using Collagenase (C8176, Sigma Aldrich) in Trypsin solution. 7-AAD cell viability dye was used to exclude non-viable cells during FACS. Single cells were sorted into lysis buffer dispensed in a 96 well plate.

### Single cell library preparation and sequencing

Single cells were processed following the Smart-seq2 protocol (Picelli et al., 2013) to reverse transcribe poly-adenylated RNA. The amount of synthesized cDNA was quantified using the Quant-iT PicoGreen dsDNA Assay (ThermoFisher). cDNA quality was confirmed on an Agilent 2100 Bioanalyzer (Agilent Technologies). Final sequencing libraries were prepared using the Nextera XT DNA Library Preparation Kit (Illumina).

### Sequencing data analysis

Transcriptome data was mapped to the zebrafish reference genome (GRCz10) using the STAR gapped aligner (Dobin et al., 2013). Duplicate reads were removed and reads were summarized using FeatureCounts (Liao et al., 2014). Quality control of the single cell read counts was done using the Scater R package (McCarthy et al., 2017). Libraries that had a size more than 3 median absolute deviations (MADs) below the median of the whole dataset were excluded from the analysis. Furthermore, libraries were excluded from the analysis if the number of expressed genes was more than 3 MADs below the median of the whole dataset, or if the percentage of counts representing spike in features was more than 3 MADs above the median of the whole dataset. Genes with an average expression across all cells of below 0.1 counts as well as mitochondrial genes were excluded from analysis. The cleaned dataset was further processed using Pagoda routines in the scde package (Fan et al., 2016; Kharchenko et al., 2014). Genes annotated to cell cycle related gene ontology terms were excluded from the dataset. Counts were transformed into FPKM expression values with the rpkm() command in the edgeR package (Robinson et al., 2010) and heatmaps drawn using pheatmap. The Rtsne package (Krijthe, 2015) was used to plot t-SNE representations of the dataset that were based on 2376 highly variable genes. RNA velocity was analysed using the Velocyto R package (La Manno et al., 2018) and pseudotime analysis was performed using the Monocle 2 package (Qiu et al., 2017). Raw and processed data generated in this study were submitted to GEO (accession number GSE121750).

### Transient Cas9 knockout

sgRNAs against the coding region of the target gene were designed to have a high Cas9-mediated nuclease cutting efficiency, a low number of genomic off-target sites and a target site proximal to the recognition site of a restriction enzyme. sgRNA primers (see Table S1) were annealed and the product inserted via Golden Gate cloning into a U6a mini-vector containing a tracrRNA backbone and Ds transposon sequences. sgRNA vectors were first injected individually to test their Cas9-mediated nuclease cutting efficiency. Typically, 30 pg of vector was injected into one-cell stage zygotes, together with 10 pg *Ac* mRNA and 160 pg *Cas9* mRNA. For actual gene editing experiments, two sgRNA vectors were co-injected at an amount of 30-38pg each. Control embryos were injected with two sgRNA vectors targeting *mCherry* (KO *tgm2b, sema3fb)* or *AmCyan* (KO *cxcl12a).*

### sgRNA Cas9-mediated nuclease activity efficiency test

One cell stage zygotes were injected with a single sgRNA vector, *Ac* mRNA and *Cas9* mRNA. Genomic DNA from single embryos was isolated at 1dpf and the genomic region surrounding the sgRNA target site (200-400bp) was amplified via PCR (see Table S2 for primers used). Half the volume of each PCR product was digested with a restriction enzyme that had a recognition site overlapping or adjacent to the sgRNA target site.

### Lineage tracing

For 4-hydroxytamoxifen (4-OHT) labeling, 10 hours post fertilisation (hpf) embryos were placed in embryo medium with 4-OHT added to a final concentration of 10 μM, from a 1 mM stock solution made in 100% ethanol. At 2dpf and 4dpf, embryos were placed in fresh 4-OHT and grown until 5dpf.

